# Explainable Machine Learning to Identify Patient-specific Biomarkers for Lung Cancer

**DOI:** 10.1101/2022.10.13.512119

**Authors:** Masrur Sobhan, Ananda Mohan Mondal

## Abstract

**Background:** Lung cancer is the leading cause of death compared to other cancers in the USA. The overall survival rate of lung cancer is not satisfactory even though there are cutting-edge treatment methods for cancers. Genomic profiling and biomarker gene identification of lung cancer patients may play a role in the therapeutics of lung cancer patients. The biomarker genes identified by most of the existing methods (statistical and machine learning based) belong to the whole cohort or population. That is why different people with the same disease get the same kind of treatment, but results in different outcomes in terms of success and side effects. So, the identification of biomarker genes for individual patients is very crucial for finding efficacious therapeutics leading to precision medicine.

**Methods:** In this study, we propose a pipeline to identify lung cancer class-specific and patient-specific key genes which may help formulate effective therapies for lung cancer patients. We have used expression profiles of two types of lung cancers, lung adenocarcinoma (LUAD) and lung squamous cell carcinoma (LUSC), and Healthy lung tissues to identify LUAD- and LUSC-specific (class-specific) and individual patient-specific key genes using an explainable machine learning approach, SHaphley Additive ExPlanations (SHAP). This approach provides scores for each of the genes for individual patients which tells us the attribution of each feature (gene) for each sample (patient).

**Result:** In this study, we applied two variations of SHAP - tree explainer and gradient explainer for which tree-based classifier, XGBoost, and deep learning-based classifier, convolutional neural network (CNN) were used as classification algorithms, respectively. Our results showed that the proposed approach successfully identified class-specific (LUAD, LUSC, and Healthy) and patient-specific key genes based on the SHAP scores.

**Conclusion:** This study demonstrated a pipeline to identify cohort-based and patient-specific biomarker genes by incorporating an explainable machine learning technique, SHAP. The patient-specific genes identified using SHAP scores may provide biological and clinical insights into the patient’s diagnosis.

## I. Introduction

Cancer is a disease in which some cells of the body grow uncontrollably and spread to other organs of the body. Cancer is a genetic disease that is caused by the changes in the genes which control the cell’s function, especially the growth and division of cells [1]. Three different kinds of genes are responsible for cancer - proto-oncogenes, tumor suppressor genes, and DNA repair genes [2], [3]. There are more than 100 types of cancers, but carcinomas are the most common type of cancer [1].

Lung cancer is the second most prevalent type of cancer [4], and it is the leading cause of death related to cancer in the United States [5]. There are mainly two types of lung cancer - non-small cell lung cancer (NSCLC) and small cell lung cancer (SCLC) [6]. Two subtypes of NSCLC are lung adenocarcinoma (LUAD) and lung squamous cell carcinoma (LUSC). There have been many studies where lung cancer biomarkers were identified. Some studies identified race-related biomarkers [7]–[9]. The researchers also applied various well-known and novel machine learning and deep learning techniques for feature selection and classification of lung cancers and other cancer types [9]–[11]. But they have mostly used machine learning and deep learning models as “black boxes.” Recently, researchers have been using various approaches to explain the black box models. Several methods have been proposed to make the machine learning and deep learning models explainable, including Shapley Sampling [12], Relevance Propagation [13], LIME [14], ANCHOR [15], and DeepLIFT [16]. But it is not clear how these methods are related and which method to select for a particular problem. To overcome this issue, Lundberg and Lee developed a unified framework for interpreting predictions, SHAP [17]. Recently there has been adequate work to explain the machine learning models using SHAP. Levy et al. used SHAP to discover important methylation states in different cell types and cancer subtypes [18]. In a more recent study, SHAP was used to explain the deep learning model which classified the cancer tissues using RNA-sequence data [19]. Most of the studies identified the global features using SHAP values which represent the average impacts of the genes on that model [20].

Researchers also use various statistical tools, such as DESeq2 [21], edgeR [22], or LIMMA [23] to identify biologically significant genes or differentially expressed genes (DEGs) [24]–[26] from differential gene expression (DGE) analysis by comparing patient cohort with healthy cohort. The DGE analysis helps to identify potential genes associated with the pathogenesis and prognosis of lung cancer [27]. The study [27] developed an integrated approach for identifying genes associated with pathogenesis and prognosis from four different sets of DEGs from four different cohorts of lung cancer patients and corresponding normal cohorts, which means that DEGs are cohort-dependent biomarker genes and do not reflect the patient-specific heterogeneity. A recent study [28] used DGE analysis to find African American (AA) and European American (EA) cohort-based lung cancer biomarkers where they showed that using principal component analysis (PCA), AA genes are able to distinguish the normal and tumor group of AA lung cancer cohort. But surprisingly, the same AA genes are also able to distinguish the normal and tumor group of EA lung cancer cohort. This observation suggests that this cohort-based study failed to discover biomarkers for a particular cohort. Another recent study [29] also used DGE analysis to find biomarkers for lung cancer using two sets of datasets- tumor and normal samples for non-treatment studies, and cell lines after treatment and cell lines before treatment for treatment studies. The hypothesis of this study is the up-regulated genes in non-treatment studies should be down-regulated in treatment studies and vice versa. But the authors found two different sets of Biomarkers without any common genes which implies that this cohort-based study failed to discover expected biomarker. Researchers also used genome wide association studies (GWAS) to find the biomarker. In one of the studies researchers found two key loci 15q25 and 5p15 for AA cohort [30]. Another study also found eighteen key loci including 15q25 and 5p15 [31]. From these two studies, we can conclude that these GWAS studies failed to identify cohort-based biomarkers. Researchers also used machine learning-based feature selection algorithms [32]–[34] to identify biomarkers for pan cancer classification which do not belong to any cancer cohort or any specific patient. These studies (DGE analysis, GWAS, and Machine Learning-based feature selection) are similar to the population-based studies where the aim is to find cohort-based genetic changes.. As a result, the same treatment provided to the patients with the same cancer type shows different outcomes among the patients [35]. This is because each patient has unique combination of genetic changes and specific genetic changes require specific treatments. That is why it is necessary to identify the patient-specific biomarkers, which we can accomplish by identifying local interpretable features by explaining the machine learning models. The patient-specific biomarkers can be used for targeted therapy leading to precision medicine which the earlier computational studies fail to identify.

We hypothesize that biomarker genes may express differently in different patients due to the variability of mutations of genes for which cohort-based gene therapy may not be beneficial to most of the patients. To solve this issue, identifying patient-specific biomarker genes is very crucial and it may aid in precision medicine or personalized medicine. In this study, we developed a pipeline to discover global and local NSCLC-associated genes using an explainable machine learning tool, SHAP. This study identified both class-specific and patient-specific genes based on SHAP scores by calculating global and local SHAP scores, respectively. To our knowledge, there has not been any study identifying lung cancer patient-specific genes using SHAP.

The later part of this paper is ordered as follows. The “Materials and Methods” section includes the cohort analysis, preparation of the dataset, and the methods used for the research. The “Experimental Results” section provides the outcome of the research and analysis of the results. We briefly discussed our result in the “Discussion” section. Finally, conclusions and the future scope is discussed in the “Conclusion” section.

## II. Materials and Methods

### A. Workflow of the study

The overall workflow of this study is shown in Fig. 1.

**Fig. 1.**
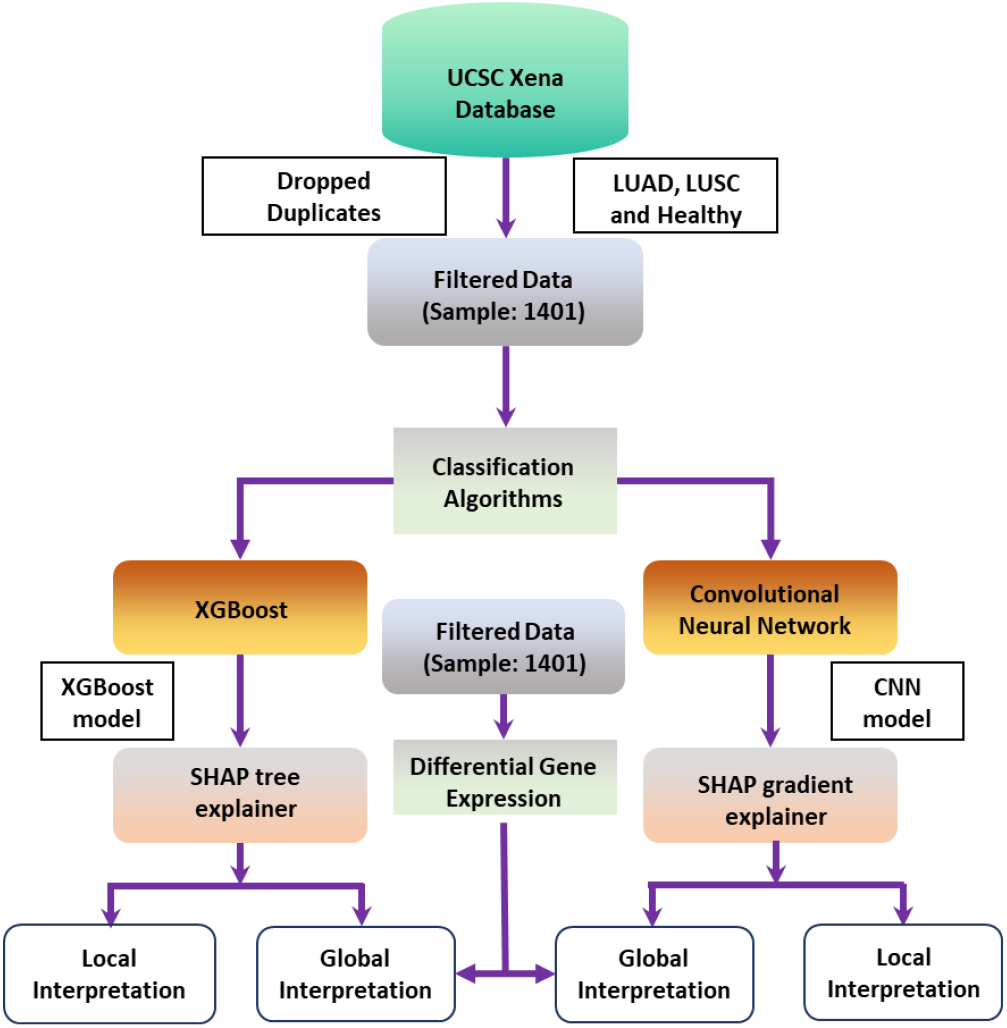
Workflow of the study to identify patient-specific and class-specific genes.

The overall workflow of this study is as follows. At first, the lung cancer tumor (LUAD and LUSC), and healthy tissue samples were downloaded from the UCSC Xena database. Next, the dataset was filtered by dropping the duplicate records of the same patients having the same tumor type. The filtered data were used to classify three different classes (LUAD, LUSC, and healthy) using two different algorithms-XGBoost and CNN. Hyperparameters were tuned to achieve a higher classification accuracy. 5-fold cross-validation was performed to measure the performance of the two algorithms. Then the two models from two different genres of classification algorithms – XGBoost from tree-based and CNN from deep learning-based classifiers - were used for interpretation using SHAP. As such, we used the tree explainer technique for the XGBoost and the gradient explainer for the CNN model for interpretation. Next, we analyzed the two different interpretation techniques to get class-specific genes and patient-specific genes. We also used a statistical tool DESEq2 to get the important genes across the populations.

### B. Data Collection and Cohort Analysis

To characterize the lung-cancer-associated mRNA, the expression profiles and clinical data associated with lung cancer were collected from the UCSC Xena database [36]. The normal tissue samples were downloaded from the Xena database and the mskcc GitHub repository [37], [38]. There are 1415 samples, including 503 LUAD, 489 LUSC, and 423 healthy, as shown in Table I.

**TABLE I.**
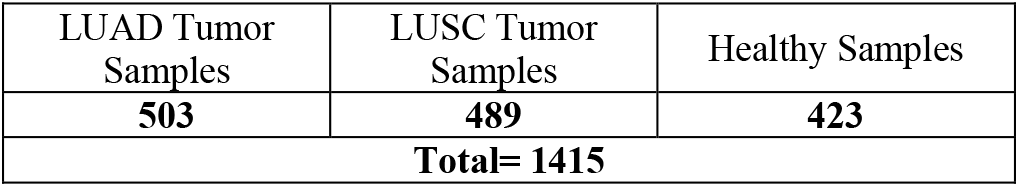
Sample distribution and cohort analysis of lung adenocarcinoma (LUAD), lung squamous cell carcinoma (LUSC) and healthy samples

| LUAD Tumor Samples | LUSC Tumor Samples | Healthy Samples |
| --- | --- | --- |
| **503** | **489** | **423** |
| **Total= 1415** |  |  |

### C. Data Preparation

Fourteen of 1415 samples were duplicates. We kept only one record of the same patients. So, the final cohort size for this analysis was 1401 with 492 LUAD tumors, 486 LUSC tumors, and 423 healthy samples, respectively. We used the dataset with FPKM values which were already log-normalized. The raw gene count dataset was also considered in this study. The data distribution of the three categories is well distributed and there is little chance of bias towards the larger group. The final dataset consists of 1401 samples with 19,648 mRNA expression values. Then we used this dataset to classify LUAD, LUSC, and healthy using a tree-based machine learning algorithm and a deep learning algorithm.

### D. Classification Algorithms

We used two algorithms in our analysis - Extreme Gradient Boosting (XGBoost) [39] and Convolutional Neural Network (CNN) [40]. XGBoost is a decision tree-based machine learning algorithm that uses a process called boosting to help improve performance. It is an optimized gradient-boosting algorithm through parallel processing, tree pruning, handling missing values, and regularization to avoid bias or overfitting.

CNN is a deep neural network primarily used in image classification or computer vision applications. But it has also wide applications in analyzing tabular data. The convolution layers extract features from the samples. A small filter or kernel scans through the samples and extracts features from the samples. The following layer is the pooling layer which down-samples the feature map extracted by the convolution layer. The pooling layer runs a filter across the feature map and takes the specific information from that filter. It translates the features’ exact spatial information to latent information. The final pooling layer is then flattened out and transformed into a one-dimensional array and fed to the fully connected layers that predict the output.

The samples of each class were divided into 80/20 split in a stratified manner for training and testing respectively. 5-fold cross-validation was used for measuring the classification performance. For the stratification of the samples, we used StratifiedKFold from the scikit-learn library. Hyperparameters were also tuned to get optimized results from both XGBoost and CNN classifiers.

Next, the contribution of all the features of individual samples for the two classifiers was determined. We wanted to identify the reasons for the machine learning models’ success or accuracy. Feature contributions, both globally and locally, can decipher the models’ accomplishment. That is why we applied an explainable machine learning tool that can identify the feature contribution that caused the models’ success.

### E. Global and Local Feature Interpretation

#### Global Feature Interpretation

The global features are a set of features that reflect the average behavior of a cohort of samples or patients. For global feature interpretation, we used two techniques: (a) DESeq2, a statistical tool, and (b) SHAP (SHapley Additive exPlanations) a game theoretic approach. DESeq2 is a tool for differential gene expression analysis of RNA-seq data. It provides a list of important genes for a cohort of patients, which reflects the average or global impact of genes across the cohort.

SHAP is a game theoretic approach to explain the output of any machine learning model. It takes the machine learning or deep learning algorithms into account and then calculates a score for each feature. The first step to calculate the SHAP score is taking the differences in the model’s prediction with and without a feature from all the coalition sets. Then taking the average of all the values from each of the coalition sets provides the SHAP score. In short, the average marginal contribution of a feature value across all possible coalitions is the SHAP score. The collective SHAP values can show how much each predictor or feature contributes, either positively or negatively, to the target variable or output of the model. The collective SHAP values refer to the global features of the dataset.

#### Local Feature Interpretation

Local features are a set of features that reflect the characteristics or behavior of an individual sample or patient. Along with identifying global important features, SHAP identifies local important features as well. Each sample for each feature or predictor gets its SHAP value. It increases transparency by calculating the contributions of the predictors. Traditional feature importance or selection algorithms produce results across the entire population, not on each individual. The idea of local interpretability of SHAP was used for identifying patient-specific genes which may help devise the strategy for personalized treatment.

## III. EXPERIMENTAL RESULT

### A. Classification Accuracy

The dataset was divided into 80/20 split for training and testing. Also, 5-fold cross-validation was performed to measure the performance of the models. The testing accuracy of algorithms from 5 folds was measured and then the average was calculated to finalize the accuracy. Table II summarizes the results of 5-fold cross-validation. The testing accuracy of XGBoost and CNN were 96.3% and 92.6%, respectively.

**TABLE II.** Results of 5-Fold Cross-Validation. First Row: Distribtuiion of Actual Labeled data; Second row: Distribtuiion of correctly predicted data; Third row: average classification accuracy of 5-fold cross-validation

|  | XGBoost |  |  | CNN |  |  |
| --- | --- | --- | --- | --- | --- | --- |
|  | LUAD | LUSC | Healthy | LUAD | LUSC | Healthy |
| Actual Data | 492 | 486 | 423 | 492 | 486 | 423 |
| Correct Prediction | 473 | 453 | 423 | 468 | 450 | 379 |
| Accuracy | 96.3% |  |  | 92.6% |  |  |

### B. Differential Gene Expression Analysis

Differential gene expression (DGE) analysis was performed using the statistical tool DESEq2. Raw counts of gene expression value were used in this analysis. We used 492 LUAD tumor samples, 486 LUSC tumor samples, 59 healthy tissues from LUAD patients, and 51 healthy tissues from LUSC patients. We ran the DGE analysis tool on LUAD and LUSC samples separately to get the most important lung cancer subtype (LUAD and LUSC) specific genes. These genes represent the average behavior of the population related to the subtypes (LUAD and LUSC). These genes can also be named global features as they represented the average importance of the cohorts. We identified LUAD-specific differentially expressed genes (LUAD-DEGs) and LUSC-specific differentially expressed genes (LUSC_DEGs) based on the thresholds |log2Fold-change| >3 and adjusted p-value < 0.001, which provided us 1,037 and 1,773 genes, respectively.

### C. Global Interpretability using SHAP

We used the explainable machine learning tool, SHAP, to identify the important genes by leveraging XGBoost and CNN classifier models. The important genes were compared with the differential gene expression genes derived from the DESeq2 tool discussed in section ‘B’. SHAP and DESeq2 tools both were used to identify the important genes across the population.

In our analysis total number of features (genes) used for XGBoost and CNN algorithms was 19,648. SHAP generates a shapely score for each gene for each patient. The scores were then averaged across the samples of correctly classified classes. Thus, we got three sets of genes (LUAD-specific, LUSC-specific, and healthy-specific) with scores. We sorted the genes of each class based on the shapely values. Both XGBoost and CNN generated 5 different models because of five-fold validation. For global interpretation, we considered the average of the five models’ output (five sets of test data from 5-fold) from XGBoost and CNN. Next, we took the top 1037 genes from LUAD and 1773 genes from LUSC class each, the same as the number of DEGs. The top genes of LUAD and LUSC classes were compared with LUAD-DEGs and LUSC-DEGs, respectively. From the analysis we noticed that the tree explainer leveraging the XGBoost model and gradient explainer for the CNN classifier model were able to identify a significant number of global genes for both LUAD and LUSC classes which are shown in Fig. 2. From the figure it is clear that XGBoost model identified 89 LUAD and 214 LUSC common genes with LUAD DEGs and LUSC DEGs respectively. Whereas CNN only identifies 68 LUAD common and 218 LUSC common genes.

**Fig. 2.**
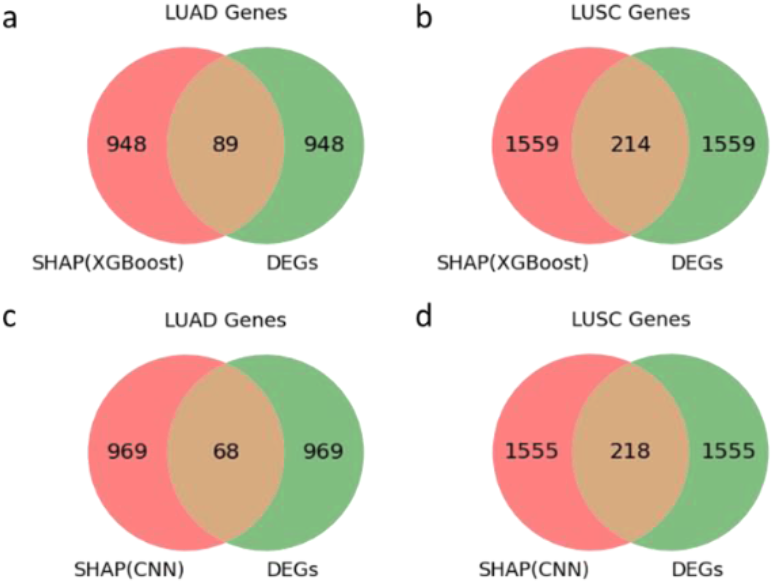
Venn diagram of SHAP genes and DEGs. (a) and (c) represent the SHAP genes and DEGs for LUAD tumor. (b) and (d) represent the SHAP genes and DEGs for LUSC tumor.

#### Optimal Genes for global interpretation

To find the optimal number of genes for global interpretation, we ran four classifiers-three variants of SVM (linear, rbf and polynomial) and logistic regression with different set of top genes. To identify the top genes, at first, the genes were sorted in a descending order based on SHAP score and then picked up the important genes. Genes having higher SHAP score are considered as the important genes. For example, top 25 genes indicate the most important 25 genes from each of the classes (LUAD, LUSC and Healthy). The criteria to select optimal number of genes was to find a minimal number of genes with high accuracy. We found out that SVM rbf and SVM polynomial are not good classifiers for the three classes. Logistic regression and SVM linear were good at classifying the three classes using the top genes. But unfortunately, SVM linear failed to classify using top 25 genes. Logistic regression and SVM linear showed that the classification accuracies were high using top 50 genes. Thus, for this study we chose top 50 genes as the optimal number of genes for global interpretation. This scenario is shown in Fig. 3.

**Fig. 3.**
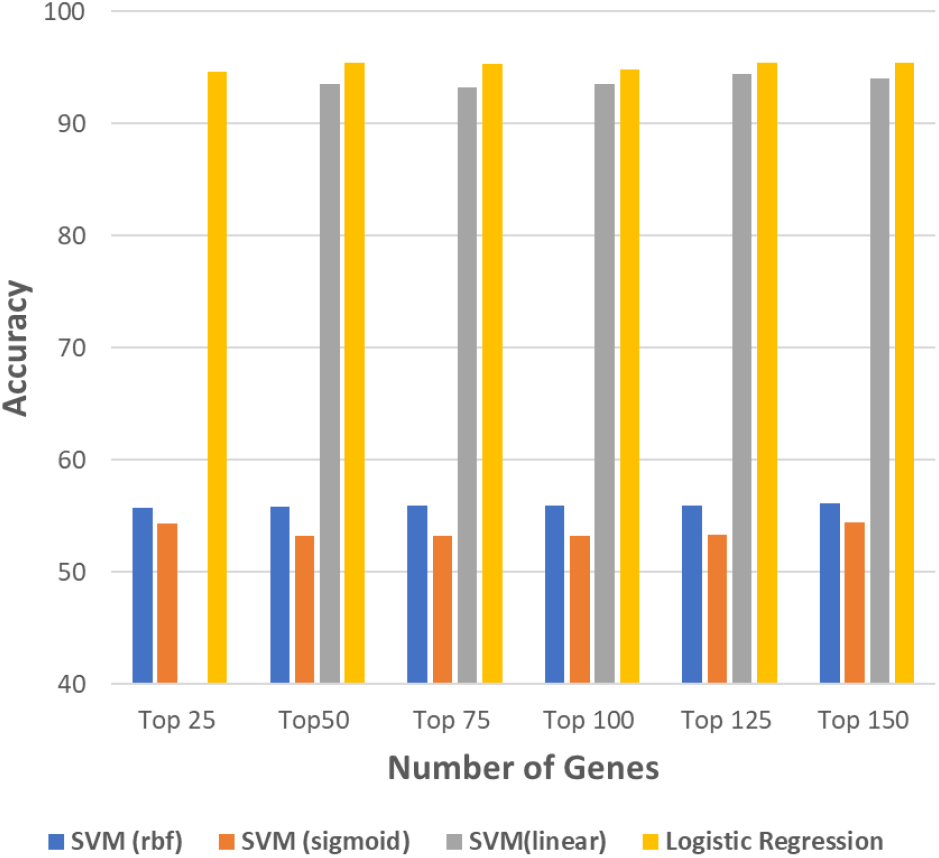
Classification accuracy of four classifiers using top genes to find optimal set of genes for global interpretation. Top 50 genes from each classes (LUAD, LUSC and Healthy) are the optimal number of genes for global interpretation as top 50 genes has a minimum number of genes with high accuracy.

Next, we examined whether the top 50 genes are truly class-specific genes. If these genes are truly class-specific then there must be few overlaps among the three groups (LUAD, LUSC, and Healthy). This scenario is shown in Fig. 4 (a). We considered the top 50 genes from LUAD, LUSC, and Healthy samples separately. We found that there is no common gene among the three sets derived from both XGBoost and CNN. There are very few or no common genes when considering two of the three classes. Also, t-SNE plot shows that, using the top 50 genes from three classes, there are three clusters for the three different classes shown in Fig. 4(b). Thus, we can conclude that the identified top 50 genes for three classes are truly class-specific. Fig. 4. only represents the genes identified by tree explainer. Similar scenarios were observed for the genes identified by gradient explainer.

**Fig. 4.**
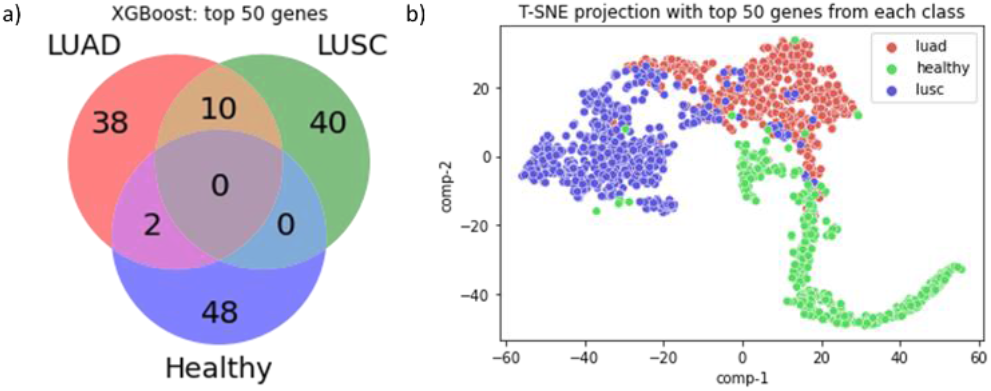
Venn diagram and t-SNE plot of class-specific genes. (a) Top 50 genes of LUAD, LUSC and Healthy from tree-explainer shows mimum overlap among the genes (b) Top 50 genes of LUAD, LUSC and Healthy from tree-explainer shows three clusters for three classes.

SHAP also provides us the information on important genes that contributed most to the model along with its shapely scores and class impact of the genes. Fig. 5 shows the top 10 genes that contributed most to the XGBoost model (CNN is not shown). It also provides information about the class-specific impact of the genes. For example, ACVRL1 (gene) contributed most to both healthy class and model output. TP63 contributed most to the LUSC class and slightly contributed to LUAD class. This means that the TP63 gene could be an important biomarker for LUSC. Similarly, we can say that GOLM1 is an important biomarker gene for LUAD.

**Fig. 5.**
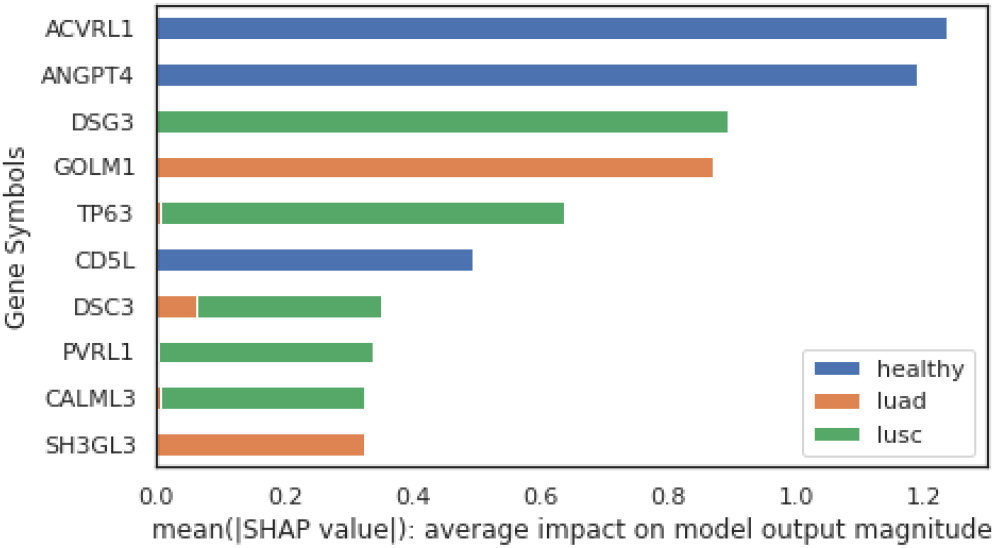
Barplot of top 10 genes. The X-axis is the mean SHAP values scored by XGBoost. The values indicate the average score of the model output for the genes. Blue, orange, and green color represent three different classes-healthy, luad, and lusc. The Y-axis represents the top gene symbols determined by the tree explainer.

### D. Local Interpretability using SHAP

We identified the most salient genes from the XGBoost and Convolutional neural network (CNN) model using SHAP for each gene and each sample. This level of local interpretability helped to identify patient-specific biomarkers which may be used as personalized medicine or therapy. To get the scores for each of the genes and samples, we trained both XGBoost and CNN with 80% of the data and tested with the rest 20% of the data. We followed this procedure five times and in each of the cases, there was a new 20% of the data in the testing set, thus providing 100% of data after testing. But XGBoost and CNN have 96.3% and 92.6% accuracy respectively which indicates that there are few false predictions. Next, we discarded the false predicted samples and kept only the true prediction. Out of 1,401 samples, the numbers of correctly classified samples were 1,349 and 1,297 for XGBoost and CNN, respectively. So, each of the samples has all the genes scored based on shapely values. Next, we sorted all the genes in descending order based on the score.

Fig. 6 shows the most important genes for a single patient. This figure is a force plot for a particular LUSC tumor patient. The predicted SHAP score of this sample is 6.23 where the base value is 0.8992. This score indicates that the expression values of the genes for this patient have a higher influence on the model. The base value is the average of the model output of LUSC class. The red arrow indicates that the genes pushed the model score higher and the blue arrow indicates the genes that pushed the model score lower. From the gene expression values, we also see that DSG3 has a high expression value and SLC4A4 has a low expression value thus the former is red and the latter is blue.

**Fig. 6.**
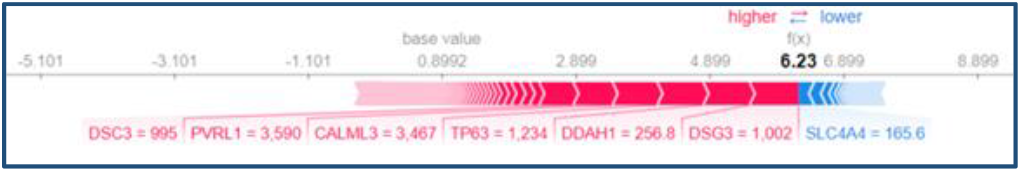
Force plot of a single LUSC patient. The numerical values along with the genes are the expression values for this patient. This plot shows the most important genes for this particular patient.

Next, we tried to interpret the patient-specific genes of each of the samples. We wanted to make sure whether these genes are really patient-specific or not. To prove it we considered randomly chosen five LUAD and five LUSC samples. For each of the patients, we picked the top 100 genes based on the SHAP score (higher SHAP-scored genes were chosen). Our hypothesis was that if these genes are really patient-specific then there should be very few overlapping genes as each individual has different mutations of genes and different expression profiles. To validate this hypothesis, we plotted a heatmap for LUAD samples which is shown in Fig. 7. From the figure, we see that there are very few overlapping genes from the tree explainer output leveraging the XGBoost model. On the other hand, the gradient explainer was able to find totally unique genes or almost zero overlapping genes among the five patients. The same scenario was observed with the LUSC patients as well (not shown). Also, the heatmap was plotted across all the patients and very few overlapping genes were found. This indicates that, even though these samples are coming from the same class, SHAP was able to score patient-specific genes.

**Fig. 7.**
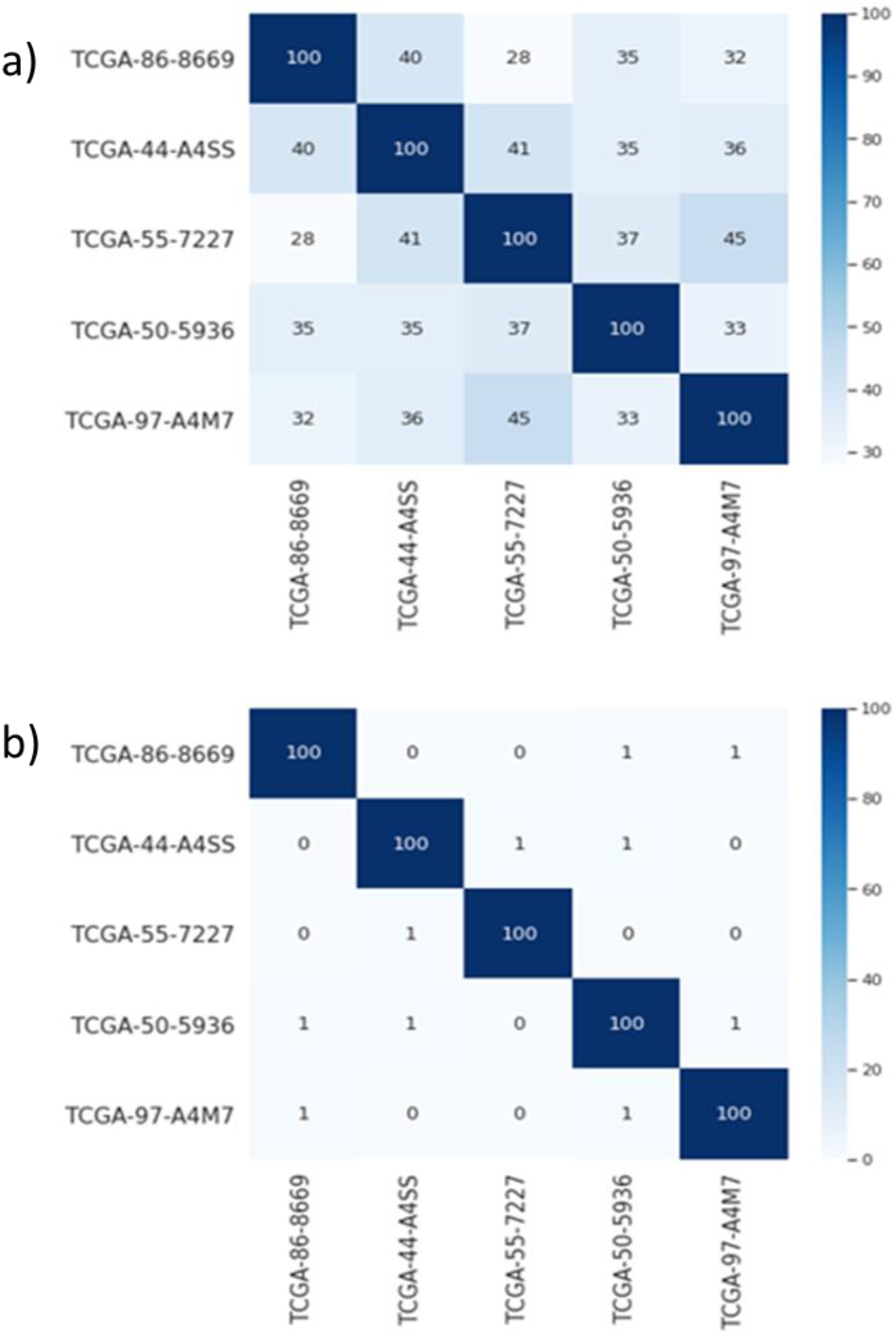
Heatmap of LUAD patients with corresponding top 100 genes. (a) Heatmap of 100 genes derived from tree explainer (XGBoost model) across 5 LUAD patients. (b) Heatmap of 100 genes derived from gradient explainer (CNN model) across 5 LUAD patients.

Next, we hypothesized that there should be many overlapping genes in the healthy samples. This is because there should be very few mutations of genes as the tissue samples are not affected by the tumor. Again, we plotted a heatmap with randomly chosen 5 healthy patients shown in Fig. 8. From the heatmap, it is evident that there are lots of overlapping genes which proves our hypothesis.

**Fig. 8.**
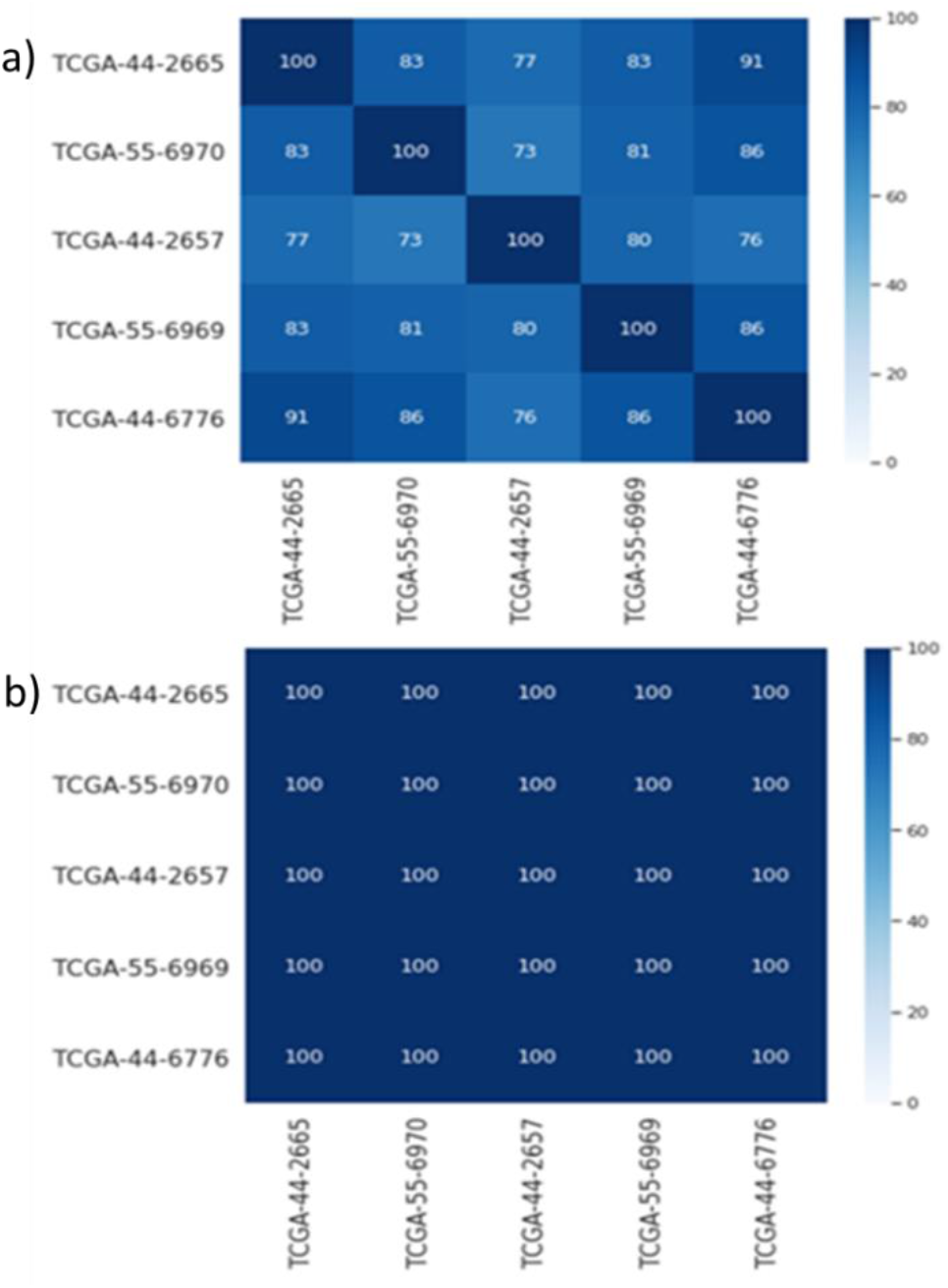
Heatmap of healthy samples with corresponding top 100 genes. (a) Heatmap of 100 genes derived from tree explainer (XGBoost model) across 5 healthy samples. (b) Heatmap of 100 genes derived from gradient explainer (CNN model) across 5 healthy samples.

## IV. Discussion

Most of the prior machine learning and deep learning works were involved in cancer classification and the algorithms were used as a “black box.” But recently a few algorithms like-SHAP, LIME, ANCHOR, DeepLIFT, etc. algorithms have been introduced to explain the black box. In this study, we used a tree-based algorithm, XGBoost, and a deep learning classifier CNN to classify the two types of lung cancer (LUAD and LUSC) and Healthy cohorts. Then the models generated by the classifiers were used in SHAP to explain the output of the models. SHAP is a unified approach to explaining machine learning models which addresses the limitations of the black box models by explaining local and global features. We used two different explainers-a tree explainer for the XGBoost model and a gradient explainer for the CNN model. Tree explainer is a fast and exact method to estimate SHAP values for the tree models. Gradient explainer is another kind of SHAP explainer that can handle neural network models. In this study, we tried to address an important task that may play a vital role in the field of healthcare, personalized medicine, by adopting the proposed pipeline.

SHAP is able to identify global features that explain the impact of the model output on the whole population. To identify whether the SHAP explainability model was able to identify plausible features, we compared the output of the two explainers with the differential gene expression (DGE) analysis tool DESEq2 output. The DGE tool was used as the reference to assess the correctness of the predicted genes from the explainers. Unlike DESEq2, there is no standard approach for selecting SHAP features (genes). That is why we ranked the genes based on the SHAP values and considered the only top-ranked genes to compare with differentially expressed genes (DEGs). The outputs of both the explainers had some common genes with the output DGE analysis. We also tried to find out the common genes among the three classes (LUAD, LUSC, and healthy) and found very few genes overlapping among the two classes, and none of the genes overlapped among the three classes. It tells us that SHAP was able to identify biologically significant class-specific genes.

One of the greatest challenges in healthcare is to identify patient-specific important biomarkers which can aid in personalized medicine. In this study, we addressed this issue by explaining the local interpretability of SHAP output. SHAP scores were assigned to every gene of every sample leveraging the modification of the game theoretic approach. So, each of the genes of every sample consists of a SHAP score which is then ranked based on the score. To explain the local interpretability, we considered the top 100 genes of each patient. We tried to find out the common genes among the samples of the same classes and found that tree explainer output has very few common genes across the samples, whereas gradient explainer has almost zero overlapping genes across the samples. It tells us that SHAP can identify patient-specific important genes in the tumor classes (LUAD and LUSC) as the tumor is more likely to work differently in different patients. We also noticed that there are lots of overlapping genes across the healthy samples. It is understandable because there is no mutation or few genomic alterations in the patients.

## V. Conclusion

Majority of previous studies identified only cohort-based important genes or population-based important genes. But it was observed that different patients require different kinds of treatment for the same disease due to the various genomic alterations and mutations. In this study, we addressed two important issues of therapeutics- the identification of subtype-specific (class-specific) and patient-specific genes. To solve these issues, we developed a pipeline that can identify both subtype-specific and patient-specific genes leveraging SHAP scores. For this analysis, we used RNA-seq data of lung cancer to show that SHAP was able to identify both class-specific and patient-specific genes. This study shows that SHAP can be used to find many biological insights by identifying local (patient-specific) and global (class-specific) genes which may help to develop better therapeutics for individual patients.

All the output shown in this analysis is machine learning and deep learning-based computational outcome. These outcomes should be verified in the wet lab to strongly validate our result. If they can be verified in the wet lab, the pipeline can be used to identify important genes for any type of disease.

## Acknowledgment

The work has been supported by the NSF CAREER (#1901628) to AMM.

